# Requirement for the Sts1 Adaptor in Proteasome Nuclear Import Elucidated in a DNA Origami NPC Mimic

**DOI:** 10.64898/2026.11.25.768562

**Authors:** Chetna Dhembla, Qingzhou Feng, Chenxiang Lin, Mark Hochstrasser

**Author notes:** Authors contributed equally (Chetna Dhembla, Qingzhou Feng).

## Abstract

The 26S proteasome, which is essential for eukaryotic protein homeostasis, comprises a 19S regulatory particle (RP) and 20S core particle (CP). Most proteasomes localize to the nucleus in exponentially growing cells, but their import mechanisms remain poorly understood. We have employed a nuclear pore complex (NPC)-mimic system to investigate translocation of the yeast proteasome *in vitro*. The system is built from DNA origami-based channels (“NuPODs”) containing densely grafted Nsp1, a Phe-Gly (FG)-rich nucleoporin. FG-repeat-containing nucleoporins create a phase-separated barrier limiting translocation of large macromolecules across the NPC. The *in vitro* system measures enhanced localization of proteasomes to NuPODs. We demonstrate that the yeast import adaptor Sts1, which binds proteasomes and provides a nuclear localization signal (NLS) for interaction with the nuclear transport receptor complex importin-α/importin-β, is required for selective association of proteasomes with NuPODs. The RP is sufficient for such NuPOD interactions, while the CP is not. Without Sts1, importin-α/β can interact with NuPODs but does not bridge proteasomes to the channels, indicating that intrinsic proteasome NLSs are insufficient for proteasome import.

## INTRODUCTION

Protein homeostasis in eukaryotic cells relies heavily on the ubiquitin-proteasome system (UPS). The 26S proteasome, a ∼2.5 MDa ATP-dependent proteolytic complex, degrades proteins tagged with ubiquitin as well as those that are targeted by other means (Finley *et al*, 2016) (Kumar Deshmukh *et al*, 2019) (Mao, 2021). The 26S proteasome comprises a 20S core particle (CP) and a 19S regulatory particle (RP) attached at one or both ends of the CP (Sahara *et al*, 2014) (Budenholzer *et al*, 2017). The CP encloses the protease active centers for protein substrate breakdown, while the RP is responsible for ubiquitin recognition, substrate unfolding, ubiquitin tag removal, and substrate channeling into the CP core (Finley, 2009) (Bard *et al*, 2018).

In exponentially growing yeast cells, ∼80% of proteasomes concentrate in the nucleus (Pack *et al*, 2014) (Wendler & Enenkel, 2019). By contrast, in nitrogen-starved or glucose-starved cells or in quiescent cells, most proteasomes relocalize to the cytoplasm. In quiescent or glucose-limited cells, proteasomes form large cytoplasmic structures called proteasome storage granules (PSGs), while under nitrogen starvation, they are degraded in the vacuole (Laporte *et al*, 2008) (Li & Hochstrasser, 2020) (Tang *et al*, 2026). Upon glucose refeeding, PSGs rapidly dissipate, and proteasomes re-enter the nucleus within minutes (Laporte *et al*., 2008). Such proteasome reimport appears to be mechanistically distinct from import of nascent proteasomes assembled in the cytoplasm (Breckel *et al*, 2024) (Weberruss *et al*, 2013).

Transport of virtually all macromolecules across the NE occurs through the nuclear pore complex (NPC); for proteins larger than ∼40 kDa, efficient import depends on specific sequence or structural elements, known as nuclear localization signals (NLSs), in the substrate (Wente & Rout, 2010). NLS-containing substrates bind nuclear transport receptors (NTRs) that ferry them across the NPC (Tetenbaum-Novatt & Rout, 2010). The NPC is composed of roughly 30 different nucleoporins (Nups), among which the phenylalanine-glycine (FG) repeat-containing Nups form a phase-separated selectivity barrier that enables specific, receptor-mediated translocation of cargo (Aitchison & Rout, 2012) (Gorlich, 2025). Cargo transport depends on transient interactions between the Nup FG repeats in the NPC channel and hydrophobic pockets on NTRs such as importin-β (Chen *et al*, 2025) (Wing & Chook, 2026).

While the structure, assembly, and biochemical functions of the proteasome have been extensively characterized, the mechanisms governing its transport across the nuclear envelope (NE), as well as the regulation of proteasome localization, remain poorly understood (Chen & Madura, 2014; Wendler & Enenkel, 2019) (Wendler & Enenkel, 2019). Nuclear import of proteasomes relies on the importin-α/importin-β NTR complex (Wendler & Enenkel, 2019). Importin-α recognizes classical NLS (cNLS) sequences and binds to importin-β, which carries substrates across the NPC central transport channel (Wing & Chook, 2026). Import substrate-receptor complexes are dissociated after binding to the small GTPase RanGTP, present at high concentration inside the nucleus (Lee *et al*, 2005) (Hirano *et al*, 2017). Both the CP and RP subunits bear cNLS-related sequences that can function as import signals when fused to heterologous reporters (Lehmann *et al*, 2002) (Wendler & Enenkel, 2019). However, whether they are capable of, or required for, directing proteasome import in their native context has been more difficult to ascertain (Chen & Madura, 2014) (Budenholzer *et al*, 2020).

Another question is whether complete 26S proteasomes, separate RP and CP complexes, or assembly intermediates are favored as import substrates (Budenholzer *et al*., 2020) (Breckel *et al*., 2024) (Lehmann *et al*., 2002) (Wendler *et al*, 2004). Potentially, several import mechanisms mediate proteasome trafficking, and their relative importance could hinge on the assembly state of the proteasome and its precursors and the kinetics of their assembly in the cytoplasm compared to import rates. Notably, the RP is assembled from base and lid subcomplexes, and no functional NLS has been identified in any of the nine lid subunits (Isono *et al*, 2007) (Budenholzer *et al*., 2020). Nevertheless, the free lid can still translocate into the nucleus in a yeast mutant bearing a large base subunit deletion; notably, such lid import did not depend on importin-α, and nuclear morphology/integrity was perturbed (Isono *et al*., 2007). The RP base can also be imported without binding to the lid, but this still requires importin-α (Isono *et al*., 2007). Intriguingly, it was recently reported that misassembled base intermediates are sequestered within the nucleus; this depends on a normally dispensable NLS in one of the base ATPase subunits (Das *et al*, 2025). Nuclear sequestration of defective base subcomplexes limits their assembly into full 26S proteasomes, which may normally proceed primarily in the cytoplasm.

Besides potential direct interaction of the proteasome with NTRs, specific adaptor proteins can bridge proteasomes to the import machinery. In *Saccharomyces cerevisiae*, the essential Sts1 protein has both a noncanonical N-terminal bipartite NLS, which binds importin-α, and a predicted 6-helix bundle that associates with the 26S proteasome (Figure 1A) (Breckel *et al*., 2024) (Budenholzer *et al*., 2020) (Chen *et al*, 2011) (Tabb *et al*, 2000). Sts1-related proteins are found in a broad range of eukaryotes, although orthologous sequences have diverged considerably (Takeda *et al*, 2011) (Budenholzer *et al*., 2020). While present in bony and lobe-finned fishes, genes encoding Sts1-like proteins appear to have been lost in the tetrapod lineage of vertebrates (Breckel *et al*., 2024). The *S. pombe* fission yeast ortholog, Cut8, has been proposed to tether proteasomes to the inner nuclear membrane by a ubiquitin-dependent mechanism rather than promote their nuclear import (Takeda & Yanagida, 2005) (Takeda *et al*., 2011).

**Figure 1.**
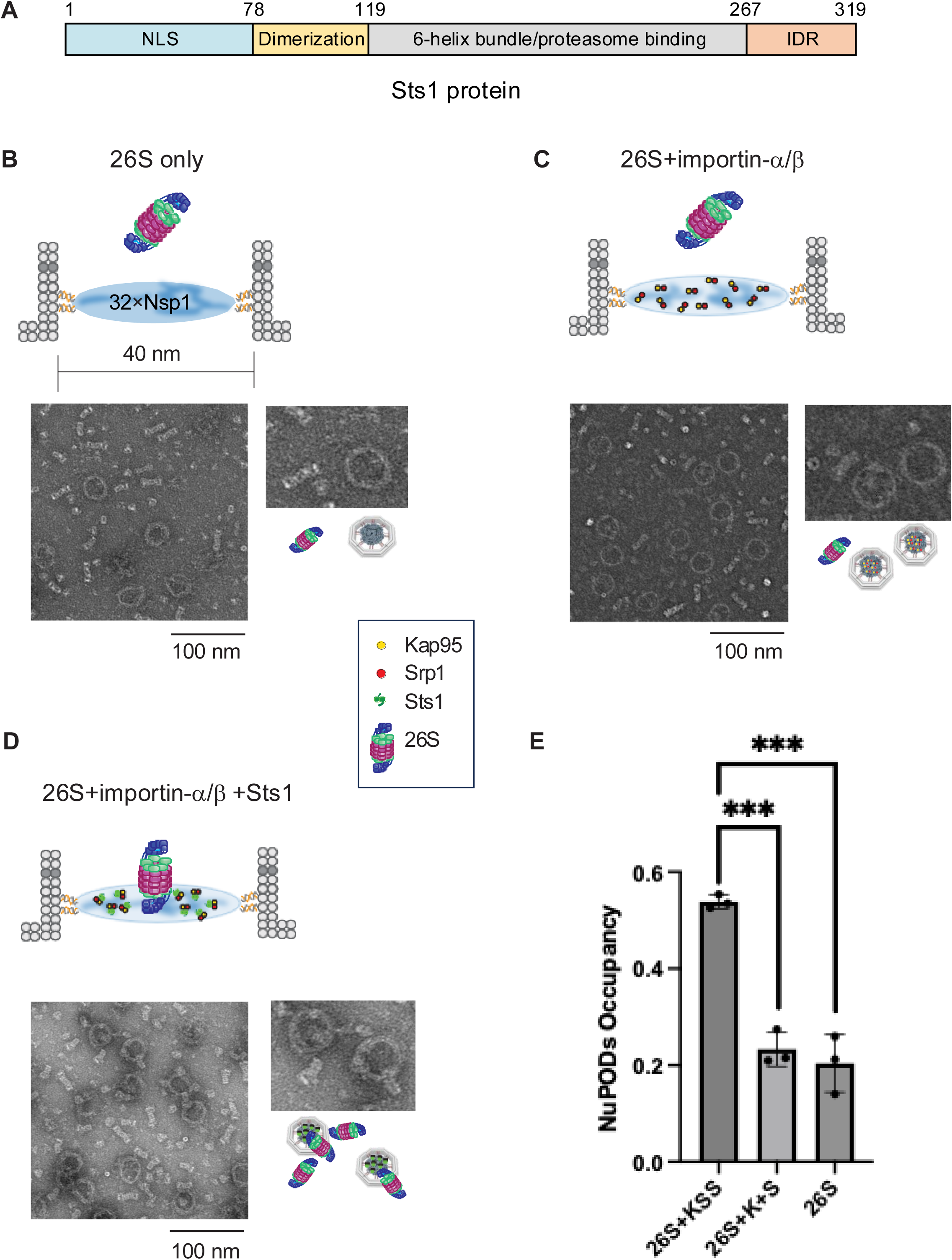
26S proteasome occupancy of NuPODs is promoted by Sts1. **(A)** Schematic of Sts1 functional domains. IDR, intrinsically disordered region. **(B-D)** TEM micrographs show the interaction of yeast 26S proteasomes with DNA nanopore-anchored nucleoporin structures (NuPODs) in the absence or presence of transport factors. Schematics of each NuPOD set-up are shown above the micrographs. Enlarged views are shown alongside schematic representations. **(B)** Incubation with 26S proteasomes alone. **(C)** 26S proteasomes added along with importins α (Srp1, S) and β (Kap95, K). **(D)** 26S proteasomes added along with importins Kap95 and Srp1 and the Sts1 adaptor (KSS complex). **(E)** Quantification of 26S proteasome occupancy of NuPODs in the presence of the indicated factors. Bar graph shows mean ± SD from three independent experiments; P-values were calculated using a two-tailed unpaired *t*-test, P values between 0.0001 and 0.001 are shown with three asterisks.

An evolutionarily unrelated protein from humans, AKIRIN2, was recently demonstrated to function in proteasome nuclear import by a mechanism with strong similarities to what has been proposed for Sts1 (Brunner *et al*, 2026; Budenholzer *et al*., 2020; de Almeida *et al*, 2021) (Budenholzer *et al*., 2020). AKIRIN2 binds directly to the CP and to several distinct importin receptors; RP import also depends on AKIRIN2, and this is likely to occur through RP piggybacking on one end of the CP barrel, with the other CP end free to interact with the adaptor (Brunner *et al*., 2026; de Almeida *et al*., 2021). AKIRIN2 is specifically required for the degradation of nuclear proteins (de Almeida *et al*., 2021). The *in cellulo* data on both yeast Sts1 and human AKIRIN2 suggest that the intrinsic NLSs of proteasomes are insufficient for their rapid nuclear import, at least for certain conditions or proteasomal assembly states such as the fully assembled 26S proteasome.

As noted above, many questions regarding the basis of proteasome nuclear import are not resolved, and they have been difficult to address in the complex environment of cells. We therefore turned to a recently described *in vitro* system for constructing NPC-like structures of defined geometry and Nup composition (Fisher *et al*, 2018) (Shen *et al*, 2023a) (Feng *et al*, 2024). Specifically, we employed NuPODs (NucleoPorins Organized by DNA Origami) composed of 40 nm-wide DNA origami scaffolds decorated with multiple copies of the yeast FG-repeat nucleoporin Nsp1. Nsp1 is an abundant central channel FG-Nup in the yeast NPC (Otto *et al*, 2024). Such nanopores recapitulate key features of the-NPC barrier and allow precise control over biochemical and biophysical parameters (Fisher *et al*., 2018; Shen *et al*, 2023b) (Shen *et al*., 2023a) (Feng *et al*., 2024). These advantages make the NuPOD system a powerful bridge between cellular observations and solution studies with soluble components (Hampoelz *et al*, 2019) (Hulsmann *et al*, 2012) (Labokha *et al*, 2013) (Shen *et al*., 2023a).

Here we have used these NPC-like nanopores to investigate the roles of the yeast import adaptor Sts1 and the classical importin-α/importin-β NTR complex in mediating nuclear import of the proteasome. Our findings highlight the critical role of Sts1 in facilitating the nuclear import of the full 26S proteasome, acting as an essential cofactor directly bridging interactions between the proteasome and the importin-α/β complex. Our data also suggest that the RP is the primary proteasome component interacting with the Sts1/importin-α/β ternary complex. The isolated CP cannot interact with the NuPODs even in the presence the Sts1-NTR complex. These insights provide a deeper understanding of the molecular mechanisms governing proteasome transport into the nucleus and underscore the importance of Sts1 as a central regulator in this process.

## RESULTS

### An *in vitro* NPC-like system to study proteasome import

It has been difficult to evaluate the direct role of Sts1 in proteasome nuclear import due to its potential involvement in other cellular processes, such as ribosome-centered quality control of nascent proteins (Ha *et al*, 2014); the presence of multiple NLSs in proteasomes that might render Sts1 redundant for import (Breckel *et al*., 2024) (Wendler & Enenkel, 2019); and the highly divergent mechanisms of action proposed for Sts1 and its *S. pombe* ortholog, Cut8, in proteasome localization (Breckel *et al*., 2024) (Chen & Madura, 2014) (Takeda & Yanagida, 2005). The components of the proteasome mediating its interaction with Sts1 also remain uncertain (Budenholzer *et al*., 2020; Chen *et al*., 2011; Tabb *et al*., 2000) (Budenholzer *et al*., 2020). We have therefore adapted the NuPOD system to address these questions with purified proteins. NuPOD structures position specific nucleoporins in defined architectures that reproduce many of the properties of NPCs, including their function as selective barriers that support NTR-mediated transport (Feng *et al*., 2024) (Shen *et al*, 2021).

Nsp1-NuPODs were assembled as described in the Methods and shown in Supplementary Figure 1. A ∼40 nm-wide octahedral channel, which approximates the symmetry and channel diameter of the NPC (Shen *et al*., 2023a), was built using DNA origami technology. The yield and geometry were confirmed by agarose gel electrophoresis and negative-stain transmission electron microscopy (TEM, Supplementary Figure 1A-E). These channels included 32 DNA oligonucleotide “handles” projecting towards the center. The handles allowed attachment of up to 32 copies of the purified yeast Nsp1 FG-Nup domains via SNAP-tag-conjugated complementary oligonucleotide “anti-handles,” mirroring the copy number of Nsp1 in the NPC central channel (Kim *et al*, 2018) (Supplementary Fig 1B,D,E). As expected for Nsp1 attachment, all NuPODs migrated more slowly than the corresponding empty DNA nanopores during electrophoresis in agarose gels (Supplementary Fig 1E, right lanes) (Fisher *et al*., 2018).

### Sts1 enhances association of 26S proteasomes with Nsp1-NuPODs

To determine whether Sts1 –as part of the importin-β (Kap95)/importin-α (Srp1) “KSS” complex is necessary for proteasome-NPC interaction, we incubated the recombinant KSS complex with 26S proteasomes purified from yeast (Supplementary Figures 2-5) and added the mixture to Nsp1-NuPODs. We also tested if, instead, the intrinsic NLSs in the 26S proteasome were sufficient for interaction with importin-α/importin-β and could thereby promote proteasome-NuPOD association without the Sts1 adaptor. Following their co-incubation, the different combinations of proteins and NuPODs were visualized by negative-stain TEM (Figure 1).

When a 4-fold molar excess of 26S proteasomes was incubated with the NuPODs in the absence of any import factors, ∼20% of NuPODs overlapped with or were occupied by proteasomes on the grid (Figure 1B, E). Addition of importin-α/importin-β (Srp1/Kap95) did not significantly alter NuPOD occupancy by proteasomes (Figure 1C, E). Remarkably, addition of the full KSS complex to proteasomes prior to NuPOD incubation led to over 50% of NuPODs associating with proteasomes (Figure 1D, E). These data strongly suggest that the importins alone were insufficient to promote 26S proteasome association with the NPC mimic and that Sts1 functions directly in linking the proteasome to the nuclear import machinery.

### Simultaneous interaction of proteasomes with NuPODs and importin adaptor complexes

It remained possible that the apparent Sts1-stimulated interaction of 26S proteasomes with NuPODs seen in Figure 1 reflected coincidental overlap of the two large complexes on the EM grids rather than stable association. To rule this out, we used rate-zonal centrifugation to characterize how the sedimentation behaviors of NuPODs, import factors, and 26S proteasomes might change when they were combined. If the KSS complex, for example, promoted stable binding between 26S proteasomes and NuPODs, we would predict a shift of the proteasomes –and KSS– to the faster-sedimenting fractions bearing Nsp1-NuPODs.

Samples were fractionated on 15–45% glycerol gradients followed by SDS-PAGE and silver staining (Figure 2). Consistent with their large molecular mass (∼9.7 MDa), NuPODs sedimented as discrete species in mid-gradient fractions; this can be seen from the ∼150 kDa Nsp1 fusion protein band that usually peaked in lanes 6 to 8 of the SDS gel (Figure 2A, B, top panels). The excess free Nsp1 protein that was left unbound during NuPOD assembly, as shown in Supplementary Figure 6A, migrated in the low-density fractions of the gradient. Notably, a fraction of importin-α/β (Srp1/Kap95) co-migrated in the higher-density fractions with the NuPODs whether Sts1 was present or not (Figure 2A, B (top panels). This shift of the NTRs was not observed unless NuPODs were present (Supplementary Fig. 6B, C) but was clearly seen in the absence of proteasomes (Supplementary Figure 7A). These results indicate that these importins can interact with the NuPODs–and by extension, NPCs–in the absence of cargo, as has been demonstrated in other experimental settings (Gorlich & Mattaj, 1996) (Cingolani *et al*, 2000) (Kozai *et al*, 2025).

**Figure 2.**
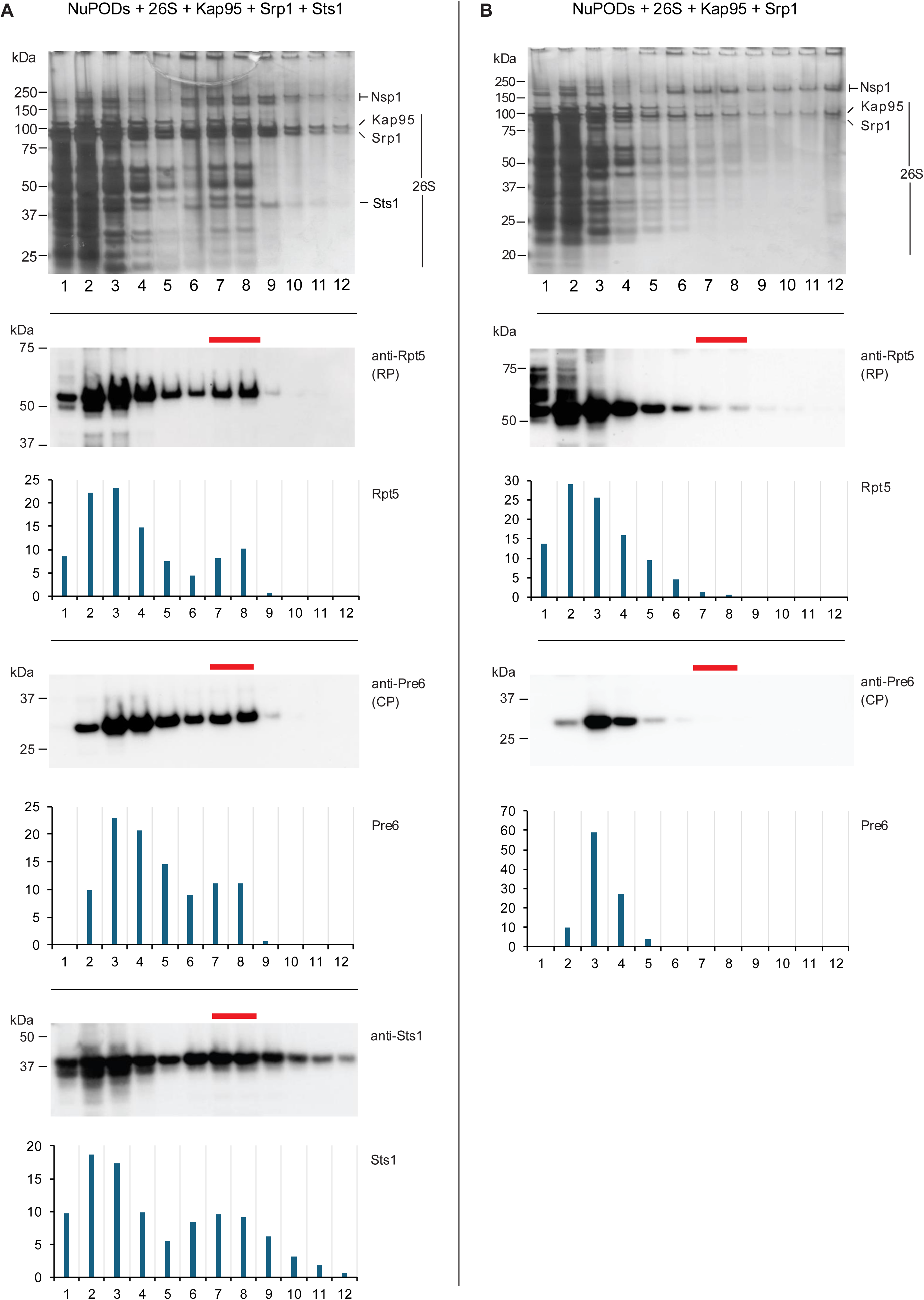
Glycerol gradient fractionation demonstrates stable Sts1-dependent interaction of 26S proteasomes with NuPODs. **(A)** In the presence of the Kap95-Srp1-Sts1 (KSS) complex, 26S proteasomes co-migrated with Nsp1-NuPODs, peaking in fractions 7 and 8 (marked by red bars). Top panel, silver staining of SDS-PAGE-separated proteins. Lower panels, Western blotting with anti-Rpt5 (RP), anti-Pre6 (CP), and anti-Sts1 antibodies confirms the shift of proteasomes to the heavier Nsp1-NuPOD fractions. **(B)** In the absence of Sts1, Kap95 and Srp1 do not promote 26S proteasome co-migration with NuPODs (fractions 7 and 8), as confirmed by immunoblot analysis with anti-Rpt5 and anti-Pre6 antibodies. Fraction numbers increase from top (low density) to bottom (high density). Molecular mass markers (kDa) are shown on the left. Blots were quantified using ImageJ; y-axis values represent the percentage of the integrated signal of the entire immunoblot in each glycerol gradient fraction.

Most importantly, when 26S proteasomes were preincubated with the three-component Sts1-NTR complex and then added to the Nsp1-NuPODs, we observed a shift of ∼15-20% of the proteasomes to the large NuPOD fractions (Figure 2A). This was observed by both silver staining of the gel and by immunoblotting with antibodies to CP (Pre6) and RP (Rpt5) proteasome subunits. The RP and CP co-sedimented, confirming the integrity of the 26S complex. Sts1, presumably as part of the KSS complex, also shifted to the denser fractions. In striking contrast, 26S proteasomes did not co-migrate with NuPODs if Sts1 was omitted (Figure 2B). As expected, this was also true if all the import factors were left out (Supplementary Figure 7B), suggesting that proteasomes do not directly engage nucleoporins in the absence of NTRs.

Several additional controls were performed. If NuPODs were omitted, neither 26S proteasomes by themselves nor proteasomes mixed with the KSS complex formed a distinct peak in the higher density fractions (Supplementary Figure 8). The ability of importin-α/β to bind NuPODs was not altered by the presence of Sts1 (Supplementary Figure 9A), and the Srp1/Sts1 complex (without Kap95) showed little if any binding to the NuPODs (Supplementary Figure 9B). In summary, interaction of 26S proteasomes with the NPC mimic required the full KSS complex.

These observations allow several conclusions. First, importin-α/β can by itself interact with the NPC mimic but does not bring the 26S proteasome into stable association with it. This implies that none of the classical NLSs that are part of both the CP and RP components of the proteasome, either by themselves or in combination, is capable of binding importin-α/β and functioning in nuclear import in this defined system. Second, inclusion of Sts1 with importins α and β is sufficient to bridge NuPOD and proteasome. In the context of the full NPC in cells, the simultaneous binding of the KSS complex to the proteasome would allow proteasomes to interact with and traverse the pore.

### The KSS complex bridges RP but not CP interaction with NuPODs

Previous protein pulldown and yeast two-hybrid results suggested that the Rpn11 subunit of the proteasome RP binds to Sts1, although the interaction was weak (Chen *et al*., 2011; Tabb *et al*., 2000). On the other hand, pulldown experiments with purified proteasomal complexes and recombinant Srp1-Sts1 suggested that the latter complex binds preferentially to the fully assembled 26S proteasome over either RP or CP (Budenholzer *et al*., 2020). To determine if either proteasomal complex engages the nuclear import machinery to allow NPC interaction, we analyzed the behavior of separately purified RP and CP in the presence of NuPODs and the KSS complex. A substantial fraction of the RP co-migrated with NuPODs in high-density fractions 7 and 8, as observed by silver staining as well as immunoblotting with anti-Rpt5 antibody (Fig. 3A). By contrast, the CP failed to co-migrate with NuPODs under identical conditions even though the KSS complex shifted with the NuPODs (Fig. 3B). Neither RP nor CP bound to NuPODs directly (Supplementary Figure 10). These results demonstrate, first, that the CP alone is insufficient for KSS-mediated NuPOD engagement and second, that the RP provides the primary interface through which the proteasome interacts with the nuclear import machinery.

**Figure 3.**
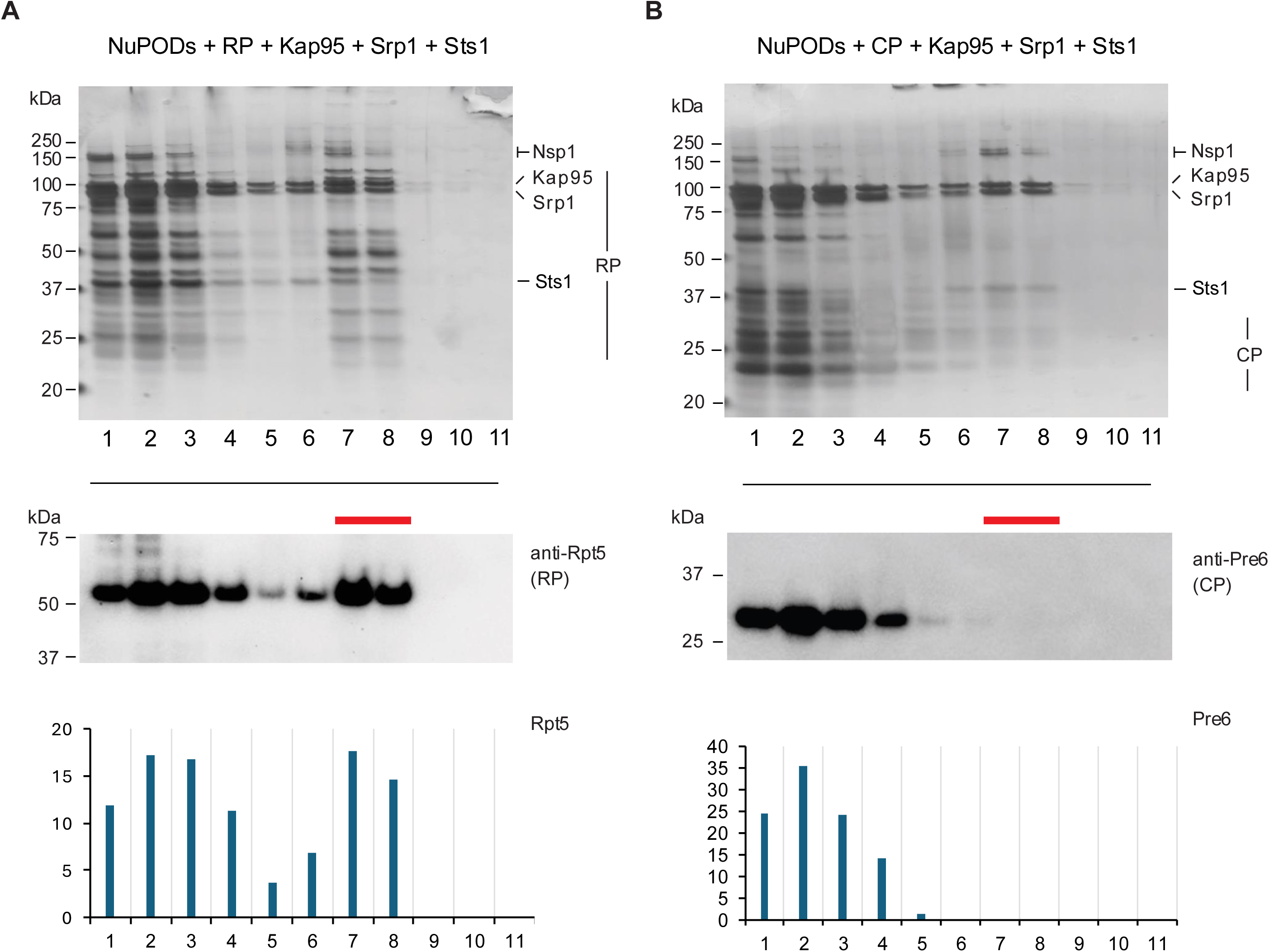
The regulatory particle (RP), but not the catalytic particle (CP), of the proteasome interacts with NuPODs. Rate-zonal centrifugation was performed to assess the interaction of individual proteasome subcomplexes with NuPODs in the presence of the adaptor complex (KSS). **(A)** The RP shifts to Nsp1-NuPOD fractions 7 and 8 (marked by red lines) in the presence of the full KSS complex. RP tracked by immunoblotting with anti-Rpt5 antibody. **(B)** The CP, when incubated with NuPODs in the presence of KSS, does not migrate to dense NuPOD fractions, indicating a lack of direct interaction under these conditions. A portion of KSS complexes did segregate to the NuPOD fractions. Fraction numbers increase from top (low density) to bottom (high density). Molecular weight markers (kDa) are shown on the left. The blots were quantified using ImageJ.

Interaction of NuPODs with RP alone could potentially be mediated directly by importin-α/β if NLS sequences within the RP become accessible when the RP is not bound to the CP. We therefore tested whether omission of Sts1 still allowed RP (or CP) interaction with NuPODs via Srp1/Kap95. As shown in Supplementary Figure 11, this was not the case: without Sts1, the RP no longer bound the Nsp1-NuPODs even though Srp1/Kap95 continued to do so. We conclude that both the full 26S proteasome and the free RP require Sts1 for importin-α/β-mediated NPC binding.

## DISCUSSION

Here we have used a previously validated, albeit simplified, *in vitro* model of the yeast NPC to investigate the nuclear transport factors required for proteasome-NPC interaction. From previous genetic and biochemical studies, the Sts1 protein had been proposed to serve as an adaptor between the importin-α/β NTR and proteasomes, driving proteasome import into the nucleus (Breckel *et al*., 2024) (Budenholzer *et al*., 2020). Sts1 has an essential N-terminal bipartite NLS sufficient for binding importin-α (Srp1). Confusingly, however, the proteasome has multiple intrinsic NLS sequences in both the CP and RP, and other research has implicated their importance for proteasome import (Wendler & Enenkel, 2019). Our results with Nsp1-NuPODs show a clear requirement for Sts1 in proteasome-NPC interaction, indicating that the intrinsic proteasomal NLSs are insufficient in the context of the full proteasome. We found that the RP by itself can also associate with NuPODs and this too requires Sts1 and importin-α/β (KSS complex). The purified CP did not bind NuPODs stably even in the presence of KSS, indicating the need for an additional factor or factors.

An updated model for Sts1-dependent nuclear import of proteasomes is shown in Figure 4. From our current data, we propose that Sts1 functions as a direct molecular bridge linking the RP of the 26S proteasome to importin-α/β. This enables multivalent interactions with FG-nucleoporins such as Nsp1 within the NPC, thereby promoting efficient docking and translocation of proteasomes through the pore. Overall, these findings are in strong agreement with *in vivo* data on the importance of Sts1 for proteasome import (Breckel *et al*., 2024; Budenholzer *et al*., 2020; Chen *et al*., 2011) (Budenholzer *et al*., 2020) (Breckel *et al*., 2024) and have refined our understanding of the process. The minimal Nsp1-NuPOD system used with purified proteasomes and import factors thus provides a valuable defined framework to evaluate mechanisms of proteasome nuclear transport. Conversely, the association of the KSS complex with NuPODs is the first example of NTR-mediated association of an NLS-bearing cargo with this NPC mimic.

**Figure 4.**
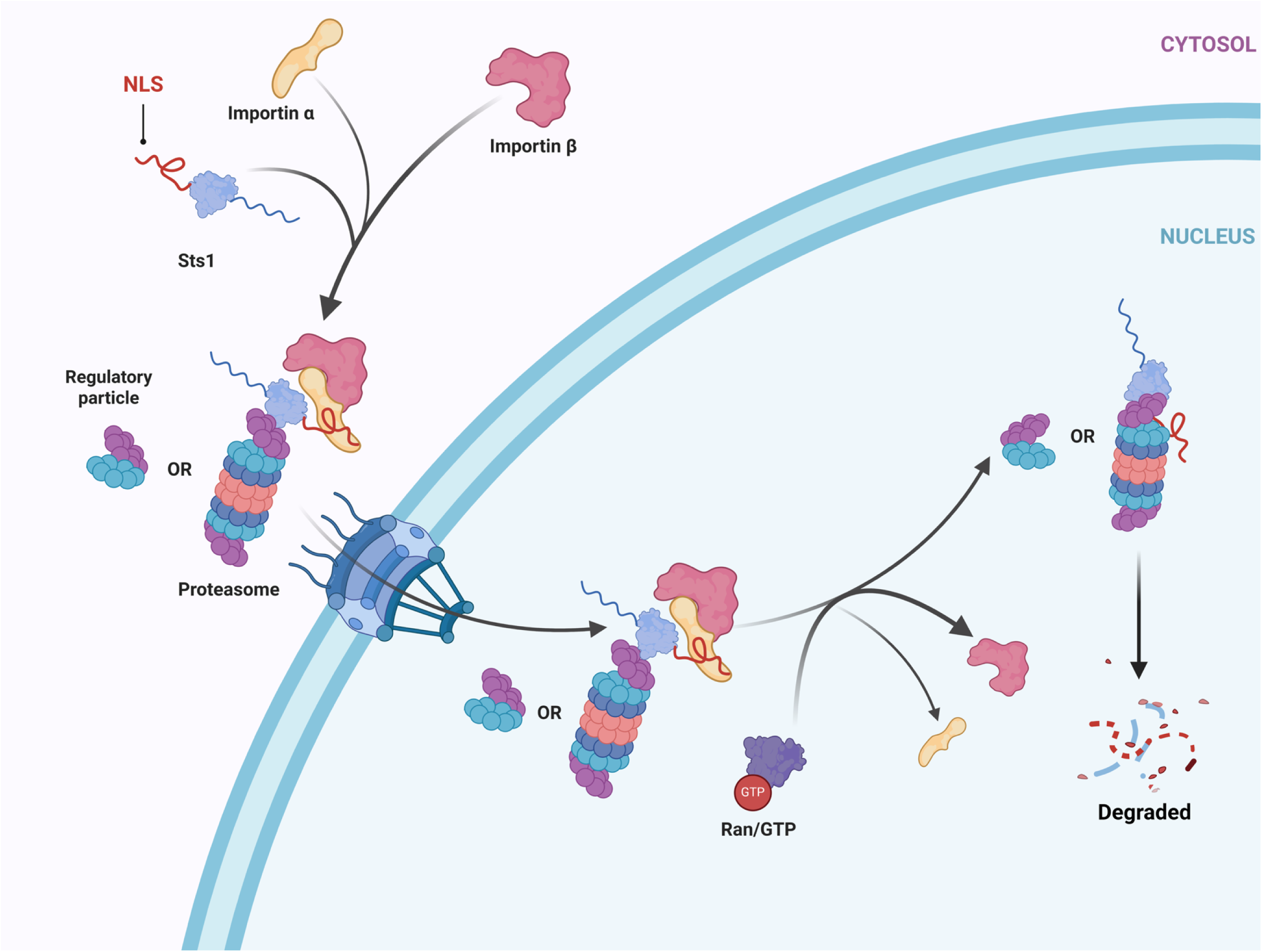
Model of Sts1 action in proteasome nuclear import. The Sts1-NTR complex appears to favor binding to the full 26S proteasome but can also bring the RP to the NPC for import. Following dissociation of the importin-α/β subunits in the nucleus, Sts1 becomes susceptible to proteasomal degradation without ubiquitin ligation, likely by the same proteasome that it helped import.

Alternative mechanisms of yeast proteasome import have been proposed (reviewed in (Wendler & Enenkel, 2019)). These distinct import routes may become more relevant under specific conditions. For instance, earlier data suggested that CP precursors are competent import substrates; our analyses with NuPODs have not addressed this possibility, which might become relevant under conditions that delay steps in cytoplasmic CP assembly (Wendler & Enenkel, 2019). Notably, following growth into stationary phase when most proteasomes accumulate in cytoplasmic PSGs, little if any Sts1 is available (Breckel *et al*., 2024) (Ohigashi *et al*, 2025), yet glucose replenishment triggers re-import of proteasomes into the nucleus within minutes. The HEAT repeat-containing proteasome regulator Blm10 appears to promote the re-import of the CP under these conditions (Weberruss *et al*., 2013). This would be congruent with the inability of the KSS complex to promote NuPOD association of purified CP (Figure 3). It will be interesting to test if addition of Blm10 instead of KSS allows CP binding to NuPODs. How the RP is re-imported under these conditions, if not associated with the CP in hybrid RP-CP-Blm10 complexes, also remains to be determined. Potentially, one or more intrinsic RP NLSs become accessible in these cells due to conformational changes or post-translational modification.

An interesting example of such a conditionally utilized intrinsic proteasome NLS was recently reported (Das *et al*., 2025). Under conditions where RP assembly is impaired, a cNLS in the Rpt2 ATPase subunit remains accessible to the importin-α/β NTR in RP assembly intermediates. As a result, these potentially aberrant intermediates are imported and sequestered in the nucleus, allowing proper proteasome assembly in the cytoplasm (Das *et al*., 2025). In the mature structure of the RP, the Rpt2 NLS is structurally occluded, but other sequences or structures that could act as NLSs might be available in cells recovering from glucose deprivation.

One apparent discrepancy between the present NuPOD data and our previous analysis of Sts1-proteasome binding is the ability of the KSS complex to promote RP-NuPOD binding, whereas our earlier pulldown experiments had suggested Sts1 preferentially interacts with full 26S proteasome complexes over either RP or CP alone (Budenholzer *et al*., 2020). This likely reflects the fact that the current analysis used a 20-fold molar excess of Sts1 (as part of KSS) over proteasome particles to allow visualization of the small and labile Sts1 protein on silver-stained gels. In our earlier pulldown experiments, we had used a 1:1 ratio of Sts1-Srp1 to proteasome. The RP is conformationally dynamic, and Sts1 might bind more tightly to specific RP conformations that occur in the context of the full 26S proteasome.

Sts1 is normally at fairly low concentrations in cells, so 26S particles fully assembled in the cytoplasm are probably the most abundant form of proteasomal particles available for import *in vivo*. Overexpression of Sts1 is sufficient to drive certain RP subcomplexes into the nucleus (Budenholzer *et al*., 2020). These observations are consistent with a role for adaptor abundance in modulating nuclear import by the nuclear import machinery and highlight the importance of considering stoichiometry when interpreting these interactions. As noted, Sts1 levels drop greatly in glucose-starved or quiescent cells, hinting at regulation of Sts1 concentration to control intracellular proteasome (or proteasome subcomplex) distribution.

In contrast to Sts1, the human AKIRIN2 import adaptor protein binds directly to the end(s) of the CP cylinder and drives import of the RP indirectly (Brunner *et al*., 2026; de Almeida *et al*., 2021). To be efficient *in vivo*, this RP import mechanism would require that a significant fraction of cytoplasmic proteasomes is only capped on one end with the RP, leaving the other end available for AKIRIN2 binding. In many mammalian tissues and cell types, there appears to be an excess of CP over RP complexes (Brooks *et al*, 2000) (Fabre *et al*, 2014) (Kim *et al*, 2025), so this requirement is likely met. By contrast, yeast cells, at least those in exponential growth, usually contain few free CP particles, and most 26S proteasomes are doubly capped (Funakoshi *et al*, 2009; Pack *et al*., 2014; Tomko & Hochstrasser, 2011). Sts1 binding to RP-CP (and RP) would be better suited to drive nuclear import under such conditions and may help explain the evolutionary retention of the Sts1 adaptor in yeast and other eukaryotes with high relative RP levels.

The defined composition of the Nsp1-NuPOD system makes clear the necessity of the Sts1 adaptor for direct interaction between the 26S proteasome and the importin-α/β receptor and the sufficiency of these components in mediating proteasome-NPC association. The system can be used to address many additional questions, such as whether, as noted above, the importin-β-related Blm10 protein can drive CP-NPC interaction. The imputed accessibility of the Rpt2 NLS in RP assembly intermediates (Das *et al*., 2025) could also be tested. Nevertheless, the Nsp1-NuPODs do not fully recapitulate the properties of native NPCs. For one, only binding, and not transport across a membrane barrier, is directly measured. For another, there are different types of FG Nups as well as distinct FG-Nup configurations within the NPC (Atkinson *et al*, 2013) (Peyro *et al*, 2021). Even with the experimental strengths of the minimalist NuPOD, these caveats are important to bear in mind. Future studies can thus take advantage of NuPOD-coupled transmembrane nanopores (Shen *et al*., 2023a) and more complex NPC mimics.

## METHODS

### Protein purification

Recombinant N-terminally glutathione S-transferase (GST)-tagged Srp1 (yeast importin-α) and C-terminally 8-histidine (8His)-tagged Sts1 were co-expressed and co-purified from *E. coli* (Budenholzer *et al*., 2020). In brief, the individual plasmids were first co-transformed into Rosetta (DE3) *E. coli* cells using pET42a(+)-based plasmids for His-tagged proteins and pGEX6P-1-based plasmids for GST-tagged proteins (see Supplementary Table 1 for plasmid list). Colonies were inoculated into LB medium containing kanamycin (50 μg/ml), ampicillin (100 μg/ml) and chloramphenicol (35 μg/ml), and the saturated overnight culture was then inoculated into 750 mL of fresh LB medium. The cultures were grown at 37°C for 5 hrs to an OD_600_ of 0.6 and induced with 0.4 mM IPTG at 18°C for 18 hrs. Bacterial cell extracts were typically produced by sonication in the presence of protease inhibitors (50 mM fresh PMSF and 1×EDTA-free protease inhibitor cocktail from Sigma), 0.1 mg/ml of lysozyme and 0.01 mg/ml DNase followed by centrifugation to clear cell debris and serial affinity purification using the respective affinity tags (Budenholzer *et al*., 2020). TALON resin (Takara) was used to enrich for the His-tagged Sts1 protein, and the eluate was then further purified on a glutathione resin (Pierce) to isolate the Sts1-His/GST-Srp1 complex. Homogeneity of eluted species was assessed by SDS-PAGE with GelCode Blue staining and by Western blotting (Supplementary Figure S4).

For purification of individual importin proteins, separate Rosetta (DE3) transformants with either a pET28a vector encoding an N-terminally His6-tagged Kap95 (yeast importin-β) (gift of C. Schlieker lab, Yale University) or pGEX6P-1 encoding GST-tagged Srp1 (Budenholzer *et al*., 2020) were used. The proteins were purified by affinity chromatography using TALON and glutathione resins, respectively. Proteins were further purified by size exclusion chromatography on a Hiload16/600 Superdex 200 pg preparative column from Cytiva (Breckel *et al*., 2024; Budenholzer *et al*., 2020) (Supplementary Figures S3, S5).

26S proteasomes were affinity purified from yeast as described previously (Li *et al*, 2023) with minor modifications (see Supplementary Table 2 for yeast strains used). Briefly, yeast cells with *RPN11* modified at its normal chromosomal locus with *3×FLAG* 3’ tag were grown in 2 L of rich medium (YPD, 4% glucose) for 24 hrs at 30°C. Cells were harvested and ground down to a fine powder in liquid nitrogen using a mortar and pestle. The resulting cell powder was flash-frozen in liquid nitrogen and stored at –80°C. For each round of purification, approximately 25 mL of cell powder were thawed and resuspended in an equal volume of lysis buffer containing 2 mM ATP. Cell lysates were clarified by centrifugation at 20,000 x g at 4°C for 20 min, and supernatants were incubated with anti-FLAG affinity resin for 2 hr at 4°C with continuous rotation. Bound proteasomes were eluted using 100 ug/ml of 3×FLAG peptide, yielding final protein concentrations of approximately 500 nM (Supplementary Figure S2). The concentration and purification of proteasomes were evaluated by SDS-PAGE and BCA protein estimation using a bovine serum albumin (BSA)-based standard curve. To purify proteasome subcomplexes, yeast expressing Rpn11-3xFLAG was used to purify the RP, and yeast expressing Pre1-3xFLAG was used to purify the CP as described previously (Li *et al*., 2023).

The sequence encoding *S. cerevisiae* Nsp1^FG^ (residues 2–603) was cloned into a pET-28a-derived vector (Novagen) flanked by His-MBP-SUMO and SNAP tag sequences and expressed in BL21-Gold (DE3) bacterial cells. Cells were grown in LB with 50 μg/mL kanamycin at 37 °C in a shaking incubator until OD_600_ reached 0.6. Nsp1^FG^ expression was induced with 1 mM IPTG at 20°C overnight. Cell pellets were collected by centrifugation and stored at −80°C until use.

For Nsp1^FG^ protein purification, cell pellets were resuspended in lysis buffer (50 mM Tris-HCl, 150 mM NaCl, 1 mM DTT, 1×Roche complete protease inhibitors, pH 7.4) and lysed in a cell disruptor (EmulsiFlex-C3, Avestin). The cell lysate was then centrifuged at 35,000 rpm for 30 min with a Type 45 Ti rotor (Beckman Coulter) to pellet cell debris. The supernatant was applied to a 0.45-µm-pore-size Millex filter before loading onto a HisTrap column (Cytiva) on an ÄKTA system (Cytiva). The column was first washed with 50 mM Tris-HCl, 150 mM NaCl, 1 mM DTT, 25 imidazole, pH 7.4, and the His-tagged protein was eluted with a 25–500 mM imidazole gradient. Peak elution fractions were further purified for protein monomers by size exclusion chromatography (SEC) using a HiLoad 16/600 Superdex 75 pg column (Cytiva).

### Assembly of Kap95-Srp1-Sts1 (KSS) complexes

Protein interaction analyses of KSS complexes were conducted as previously described with slight modifications (Breckel *et al*., 2024). Briefly, purified 7 uM recombinant bait proteins GST-Srp1and Sts1-His co-purified from *E. coli* were immobilized on 50 μL (packed volume) of glutathione beads that had been washed with 1 mL binding buffer (50 mM HEPES pH7.5, 150 mM NaCl, 1 mM MgCl_2_, 1 mM DTT). Binding of the complex to the resin was done in 2 mL of binding buffer containing fresh 1×Roche protease inhibitors (without EDTA) by rotating together at 4°C for 2 hrs. Beads were centrifuged at 4°C (1500 rpm, 2 min), washed with 1 mL of binding buffer and subsequently incubated in 2 mL of a two-fold molar excess of His-Kap95 at 4°C for 2 hrs on a rotator. Beads were then centrifuged as before and washed four times with 1 mL of binding buffer. Bound proteins were eluted from the beads with 2 mL of 10 mM glutathione in binding buffer and analyzed by SDS-PAGE and GelCode Blue (Thermo Scientific) staining. The KSS complex was further purified by gel filtration on an Superose 6 Increase column (Cytiva), and immunoblotting was done to validate the presence of all the components of the complex (Suppl. Figure S4).

### Antibodies and immunoblotting

A rabbit antibody to Sts1 was previously generated (Budenholzer *et al*., 2020). Immunoblotting was performed using the following primary antibodies: anti-GST (EPR4236, ab1 11947, Abcam; 1:5000); anti-His (MAB050, R&D Systems; 1:1000); anti-Pre6 (gift from Dr. D. Wolf lab, 1:5000) and anti-Rpt5 (BML-PW8245, Enzo Life Sciences, Batch number Z01946, 1:10000). Donkey raised anti-rabbit IgG linked to horseradish peroxidase (catalogue no NA934V, GE Healthcare; 1:10,000) or goat anti-mouse IgG linked to horseradish peroxidase (catalogue no 1031-01, Southern Biotech; 1:10,000) was used as the secondary antibody. Protein signals were detected by enhanced chemiluminescence (ECL) and quantified using a ChemiDoc system from Bio-Rad. PVDF membranes with a pore size of 0.45 micron from Merck Millipore was used for immunoblotting.

### Construction of DNA nanopore channels with yeast Nsp1 (Nsp1-NuPODs)

Origami channels were derivatized by mixing M13mp18 bacteriophage-derived circular ssDNA (8064 nt, 20 nM) with staple (including “handle”) oligonucleotides (120 nM each) and going through a 36-hr temperature gradient (85–25 °C) annealing process. The handle and origami design were described before (Shen *et al*., 2023a). The well-folded DNA nanopores were then purified from mis-folded structures and excess staples by glycerol gradient fractionation (Lin *et al*., 2013). The target fractions bearing the origami channels (Supplementary Figure S1) were collected, pooled, and buffer-exchanged with 1×TE (5 mM Tris HCl, pH 8.0, 1 mM EDTA) containing 15 mM MgCl_2_ using Amicon Ultra centrifugal tubes with a NMWL of 10 kDa (Millipore Sigma).

The purified Nsp1^FG^ fusion protein (Supplementary Figure S1F) was conjugated to ssDNA handles (5′amine-AAATTATCTACCACAACTCAC-3′,) as described (Feng *et al*., 2024; Fisher *et al*., 2018) (Shen *et al*., 2023a). Briefly, bacterially expressed Nsp1^FG^ protein was purified on a 5 mL HisTrap column (Cytiva) on an ÄKTA system (Cytiva). A single-stranded DNA “anti-handle” with a 5’ amine group was first derivatized with a BG-GLA-NHS (benzylguanine-glutaric acid-N-hydroxysuccinimide) (New England Biolabs) linker by reacting the NHS moiety with the 5’ amine group on the DNA. The BG-coupled DNA was then covalently linked to the SNAP tag on Nsp1 (Supplementary Figure 1F), with unconjugated DNA removed by SEC. By hybridizing an excess of the ssDNA (anti-handle)-conjugated Nsp1 with the origami channels bearing 32 handles (ssDNA complementary to the anti-handle), the NuPODs becomeoaded with close to 32 copies of Nsp1 FG-Nups (Ketterer *et al*, 2018) (Supplementary Figure S1).

### Rate zonal centrifugation, negative stain-TEM analysis, and quantification

Purified <u>K</u>ap95/<u>S</u>rp1/<u>S</u>ts1 (KSS), Kap95/Srp1 (KS), or Srp1/Sts1 (SS) complexes were mixed with 26S proteasomes (26S), CP (20S), or RP (19S) in buffer containing 50 mM HEPES (pH 7.5), 5 mM MgCl₂, 150 mM NaCl, 10% (v/v) glycerol, and 1 mM ATP, and incubated on ice for 30 min. NuPODs were maintained in buffer containing 50 mM Tris-HCl (pH 7.4), 15 mM MgCl₂, 150 mM NaCl, and 10% (v/v) glycerol.

The transport receptor-proteasome mixtures were then combined with the Nsp1-NuPODs, and ATP was added to maintain a final concentration of 1 mM in buffer containing 50 mM HEPES (pH 7.5), 5 mM MgCl₂, 150 mM NaCl, and 10% (v/v) glycerol. Typically, 6 nM Nsp1-NuPODs was mixed with 24 nM proteasomes (full 26S, CP, or RP) and 480 nM importin complexes (KSS, KS, or SS). The sample volume was adjusted to 100 µl, and the mixtures were incubated on ice for an additional 15–20 min prior to loading onto a 0.56 ml 15–45% glycerol (5% steps) density gradient manually made in Ultra clear 5×41mm 0.8 ml tubes (Beckman Coulter), as described previously (Lin *et al*, 2013). Samples were centrifuged for 45 min at 48,000 rpm in a Sw 55 Ti rotor at 4°C, and then fourteen equal volume 43 µl fractions were collected manually from top to bottom of the centrifuge tube using long neck gel loading tips. Aliquots (20 µl per fraction), mixed with 3× Laemmli buffer, were resolved on 4–20% NuPAGE gradient gels (Thermo Fisher Scientific). The gels were visualized by both silver staining and immunoblotting.

For transmission electron microscopy (TEM) analysis of protein complex-NuPOD interactions immediately after co-incubation, samples were spotted on 400-mesh carbon-coated copper grids and stained with 2% uranyl acetate using standard protocols (Feng *et al*., 2024). Imaging was done on a JEOL JEM-1400Plus microscope with a 4k×3k CCD camera (Advanced Microscopy Technologies). The concentrations used for NuPOD generation were 5 nM DNA origami + 320 nM Nsp1. Kap95, Srp1, KSS complexes were all used at 100 nM, while 26S proteasomes were at 10 nM. EM micrographs were analysed using RELION and Fiji software, and proteasome-NuPOD binding was quantified using GraphPad Prism. Two-tailed t-test was applied with p=0.05 as the significance cut-off.

## Data availability

This study includes no data deposited in external repositories.

## Author contributions

CD contributed to the experimental design, performed the experiments, and wrote the initial draft of the manuscript. QF assembled the NuPODs, performed the TEM analyses, helped with running the glycerol gradients and helped in drafting figures and the manuscript methods section. CL analyzed the data, contributed to the experimental design, reviewed the manuscript, and obtained funding. MH designed the study, analyzed the data, wrote the final version of the manuscript, and obtained funding. All authors read and approved the final manuscript.

## Disclosure and competing interest statement

The authors declare that no competing interests exist.

## Acknowledgements

We thank Isadora Rocha de Abreu and Zane Johnson for comments on the manuscript. The study was supported by an NIH grants R35-GM136325 to MH and R35-GM149264 and R01-AI162260 to CL.

## Notes

### Competing Interest Statement

The authors have declared no competing interest.

